# A biodegradable microgrooved and tissue mechanocompatible citrate-based scaffold improves bladder tissue regeneration

**DOI:** 10.1101/2024.02.15.580554

**Authors:** Madeleine Goedegebuure, Matthew I. Bury, Xinlong Wang, Pasquale Sanfelice, Federico Cammarata, Larry Wang, Tiffany T. Sharma, Nachiket Rajinikanth, Vikram Karra, Vidhika Siddha, Arun K. Sharma, Guillermo A. Ameer

## Abstract

Chronic bladder dysfunction due to bladder disease or trauma is detrimental to affected patients as it can lead to increased risk of upper urinary tract dysfunction. Current treatment options include surgical interventions that enlarge the bladder with autologous bowel tissue to alleviate pressure on the upper urinary tract. This highly invasive procedure, termed bladder augmentation enterocystoplasty (BAE), significantly increases the risk of patient morbidity and mortality due to the incompatibility between the bowel and bladder tissue. Therefore, patients would significantly benefit from an alternative treatment strategy that can regenerate healthy tissue and restore overall bladder function. Previous research has demonstrated the potential of citrate-based scaffolds co-seeded with bone marrow-derived stem/progenitor cells as an alternative graft for bladder augmentation. Recognizing that contact guidance can potentially influence tissue regeneration, we hypothesized that patterned scaffolds would modulate cell responses and improve overall quality of the regenerated bladder tissue. We fabricated microgrooved (MG) scaffolds using the citrate-based biomaterial poly(1,8-octamethylene-citrate-*co-*octanol) (POCO) and co-seeded them with human bone marrow-derived mesenchymal stromal cells (MSCs) and CD34^+^ hematopoietic stem/progenitor cells (HSPCs). Microgrooved POCO scaffolds supported MSC and HSPC attachment, and MSC alignment within the microgrooves. All scaffolds were characterized and assessed for bladder tissue regeneration in an established nude rat bladder augmentation model. In all cases, normal physiological function was maintained post-augmentation, even without the presence of stem/progenitor cells. Urodynamic testing at 4-weeks post-augmentation for all experimental groups demonstrated that bladder capacity increased and bladder compliance was normal. Histological evaluation of the regenerated tissue revealed that cell-seeded scaffolds restored normal bladder smooth muscle content and resulted in increased revascularization and peripheral nerve regeneration. The presence of microgrooves on the cell-seeded scaffolds increased microvasculature formation by 20% and urothelium layer thickness by 25% in the regenerating tissue. Thus, this work demonstrates that micropatterning affects bladder regeneration to improve overall anatomical structure and re-establish bladder physiology.

## Introduction

Bladder reconstruction, typically in the form of bladder augmentation enterocystoplasty (BAE), is the current standard of care for patients with severe and end-stage bladder disease.^1^ Patients can develop end-stage bladder disease from a number of conditions including interstitial cystitis, urological cancers, neurological conditions, or traumatic injuries that result in prolonged bladder dysfunction. Persistent bladder dysfunction increases pressure on the upper urinary tract and poses a critical risk to kidney damage. As such, surgical intervention via enterocystoplasty involves enlarging the bladder with autologous bowel tissue to relieve pressure on the upper urinary tract. Unfortunately, an estimated third of patients that undergo BAE experience complications due to incompatibility between the native bladder tissue and the grafted bowel tissue,^2^ and some patients will inevitably be required to self-catheterize for the remainder of their lives. The shortcomings of this procedure have prompted the development of regenerative engineering strategies to improve patient outcome. Many alternative grafts to autologous bowel tissue have been investigated, including decellularized bladder extracellular matrix (ECM), small intestine submucosa (SIS), and poly(lactic-co-glycolic) acid (PLGA).^3,4^ The most recent clinical trials for alternative bladder augmentation grafts were performed 10 years ago with very limited success in improving bladder function or supporting development of mature and functional tissue.^5–8^ Since then, a multitude of strategies have been explored to develop more sophisticated scaffolds that contain physical and biological cues to promote rapid bladder regeneration, building on the shortcomings of previous attempts.

The regeneration of vascular and peripheral nerve networks in particular is widely recognized as a major hurdle in bladder tissue engineering.^3,9^ Currently, many efforts in tissue engineering are focused on incorporating biologic factors into scaffolds to specifically promote revascularization or nerve regeneration. These strategies include immobilization of proangiogenic factors, such as vascular endothelial growth factor (VEGF) and platelet derived growth factor (PDGF), or fabricating pre-vascularized grafts with endothelial cells.^9,10^ Similarly, use of progenitor nerve cells or protein coatings to promote adhesion and neural differentiation have been suggested as alternatives for promoting development of a nerve network.^3,11,12^ All of these methods require complex graft preparation and may still be insufficient for supporting innervation and vascularization to the degree where the regenerated tissue is comparable to the native tissue. Furthermore, current strategies for revascularization and innervation require significant optimization and pose challenges for clinical translation because they rely on isolation and delivery of biologic factors to the graft.

Recent work has shown that microtopography engineering has the potential to improve or accelerate tissue regeneration, including blood vessels and nerves, through biophysical cues alone.^13–15^ Biophysical cues are known to affect cell function and overall tissue organization through contact guidance. Topography of a biomaterial can be designed to influence cell-material interactions and control cell morphology, proliferation, and differentiation.^16^ One of the most investigated topographies is microgrooves because they induce cell alignment along the major axis of the grooves.^17,18^ Furthermore, microgrooved topographies have been shown to promote endothelial cell, smooth muscle cell, and nerve cell migration.^19,20^ This effect, then, is particularly advantageous for bladder tissue engineering because guided migration of these cell types could potentially allow for efficient regeneration of the bladder wall post-augmentation, ensuring proper storage of urine and preventing scar tissue formation for overall improved recovery of function.

Given the potential of microtopography engineering to influence tissue regeneration *in vivo*,^13^ we hypothesized that a microgrooved scaffold could promote efficient and robust regeneration of the bladder wall by inducing cell alignment and migration onto the scaffold. We have shown that the citrate-based biomaterial poly(1,8-octamethylene-citrate-*co-*octanol) (POCO) co-seeded with autologous bone marrow-derived mesenchymal stromal cells (MSCs) and CD34^+^ hematopoietic stem cells (HSPCs) can support bladder regeneration, including urothelium, smooth muscle layers, blood vessels, and nerves in a baboon bladder augmentation model.^21^ In the current study, we fabricated microgrooved POCO scaffolds using a photolithography mold; characterized their mechanical properties and cell-material interactions compared to poly (1,8-octamethylene citrate) (POC) scaffolds, a previously studied biomaterial for bladder regeneration; and assessed regeneration in a nude rat bladder augmentation model. Overall, the POCO scaffolds supported normal bladder urodynamics and the quality of the regenerated tissue was significantly improved with the addition of stem/progenitor cells and microgrooves. Creating microgrooves on the scaffold specifically resulted in increased formation of neovasculature, promoted ingrowth of peripheral nerves, and increased urothelium uniformity and thickness. Thus, here we demonstrate the use of microtopography engineering as a novel, alternative approach to improve bladder tissue engineering by invoking cell contact guidance for bladder tissue regeneration.

## Materials/Methods

### Polymer synthesis

Poly (1,8-octamethylene-citrate-*co-* octanol) (POCO) was synthesized by adding octanol, 1,8-octanediol, and citric acid (Sigma Aldrich, St. Louis, MO) to a round bottom flask in a 0.2: 0.8: 1 molar ratio. The mixture was melted in a silicon oil bath at 165°C under nitrogen gas flow with stirring for about 15 min. Once melted, the flask was transferred to another oil bath at 140°C and reacted for another 3 hrs under nitrogen gas flow. The pre-polymer was dissolved in ethanol and purified by precipitation in Milli-Q water with 20% ethanol. Purification was repeated 2 more times in Milli-Q water and the final precipitation was collected and frozen at -80°C for 12-16 hrs. The pre-polymer was then lyophilized for 2-3 days until clear. The pre-polymer was dissolved in 40% ethanol w/v to form the working pre-polymer solution. POCO was characterized using proton nuclear magnetic resonance (^1^H-NMR, X500, Bruker) and mass spectrometry (AmaZon-SL, Bruker). Poly (1,8-octamethylene citrate) (POC) was synthesized as previously described.^22,23^ Briefly, 1, 8-octanediol and citric acid were mixed in a 1:1 molar ratio in a round bottom flask. The mixture was heated to 165°C under nitrogen gas flow to melt the monomers then polymerized at 140°C under nitrogen gas flow for 1 hr. POC was purified and validated following the same protocol as POCO.

### Scaffold fabrication

POCO and POC scaffolds were cured on glass slides with a poly(vinyl alcohol) (PVA) (Sigma Aldrich) sacrificial layer. 38 mm x 75 mm glass slides were cleaned with pressurized air then prepared for coating using an oxygen plasma surface treatment. After the plasma treatment, 4 mL of a 5% PVA solution was pipetted on the surface and cured at 80°C for 2-3 hrs. 1.5 mL of the working POCO or POC solution was pipetted on the PVA-glass slides and cured overnight at room temperature then 80°C for 3-4 days. After curing, slides were submerged in DI water and incubated at 60°C for 12-16 hrs to dissolve the PVA sacrificial layer and release the POC or POCO scaffolds. The functional groups of POCO and POC were confirmed with Fourier transform infrared spectroscopy (FTIR, Nicolet iS50, Thermo Scientific). The scaffolds were sterilized with an AN74i Anprolene Ethylene Oxide (EtO) Gas Sterilizer System (Cole-Parmer, Shanghai, China) for 12 hrs then leached to remove any unreacted monomer and adjust the material to physiological pH. First, the scaffolds were incubated in 100% ethanol for 30 min at 37°C then 20% ethanol in phosphate buffer saline (PBS) for 24 hrs at 37°C. The scaffolds were then leached for 7 days alternating every 24 hrs between serum-free DMEM (Thermo Fisher, Hillsboro, OR) and PBS, both with 1% Pen-strep (Thermo Fisher) and 0.1% anti-mycotic (FUNGIZONE, Thermo Fisher).

### Scaffold characterization

The tensile stress-strain properties of leached scaffolds were obtained using an Instron 5544 mechanical tester (Instron, Grove City, PA). The POCO and POC scaffolds were cut using a dog bone mold and stretched at a constant rate of 15 mm/min. The Young’s modulus and tensile strain at maximum force were measured and plotted for POC and POCO (n=3). Prism9 (GraphPad Software, Boston, MA) was used to run a Shapiro-Wilk test for normality, then perform an unpaired, two-tailed t-test to determine statistical significance. To obtain the tensile stress-strain curve of native rat bladder, tissue was isolated from a rat and incubated in PBS. Immediately post-isolation, the top and bottom of the bladder were removed to create a uniform, open loop of tissue from the middle of the bladder. Two paperclips were used to hook the ends of the tissue and load it into an Instron 5544 mechanical tester. The tissue was stretched at a constant rate of 15 mm/min, and the Young’s modulus and tensile strain at maximum force were measured and plotted.

Antioxidant activity of the leached scaffolds was tested using a DPPH assay as previously described.^24^ Briefly, 30 g punches 8 mm in diameter of POCO and POC were incubated in 0.5 mL of DPPH solution for 15, 30, and 45 min at 37°C. At the end of each time point, 150 uL of the solution was removed and absorbance for each sample was read at 517 nm using a BioTek Cytation 5 Cell Imaging Multimode Reader (Agilent Technologies, Santa Clara, CA). Final absorbance was calculated by measuring the difference between a blank and the sample and plotted as mean ± standard deviation (n=3). A Shapiro-Wilk test and 2-way ANOVA test was performed with Prism9 to calculate statistical significance (alpha= 0.05). Contact angle of the leached scaffolds was also evaluated using the VCA Optima XE (AST Products Inc. Billerica, MA). A 200 uL droplet of DI water was dispensed onto the scaffolds and contact angle was measured manually with the VCA Optima XE software. Measured contact angles for POC and POCO were plotted (n=5) and a Shapiro-Wilk test as well as an unpaired, two-tailed t-test was performed using Prism9.

Crosslinking density of POC and POCO scaffolds were calculated as previously described from the equation derived from the theory of rubber elasticity: 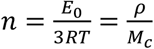 where n is the number of active network chain segments per unit volume (mol/m^3^) and M_c_ is the molecular weight between crosslinks (g/mol).^25^ 1 mm thick POC and POCO scaffolds were cured for 4-6 days at 80°C. Rectangles (0.5 cm x 2 cm) were cut from the scaffolds and the Young’s modulus (Pa) was obtained using an Instron 5544 mechanical tester as described above. 4 mm diameter punches from the same scaffolds were used to measure density. n and M_c_ were calculated from these measurements, then plotted and analyzed in Prism9 (n=4) using a Shapiro-Wilk test for normality and a two-tailed t-test.

Swelling ratio of POC and POCO was measured by comparing the wet and dry mass of the scaffolds. The samples were incubated in DI water overnight at room temperature, and the wet mass was measured the following morning. The samples were then lyophilized for 24 hrs and the dry mass was measured. Swelling ratio was calculated with the following equation: Swelling ratio 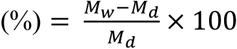. The results were plotted and analyzed using a Shapiro-Wilk test for normality and a two-tailed t-test in Prism9 (n=4).

Scaffold degradation was assessed in vitro via gravimetric analysis. POC and POCO samples 1 mm thick and 4 mm in diameter were placed in glass vials containing 3 mL of 0.1 M NaOH to obtain an accelerated degradation curve. After incubating in the NaOH for a designated time, each sample was washed 3 times with DI water and dried overnight at 65°C. Samples were weighed on an Mettler Toledo analytical balance (Thermo Fisher). The mass loss was calculated from the following equation: Mass loss 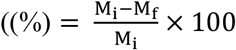. The results were plotted and analyzed using a Shapiro-Wilk test for normality and a two-tailed t-test in Prism9 (n=4).

### Microgrooved scaffold fabrication

Silicon wafers were patterned with 6-1 cm squares of 50 μm wide and 10 μm deep SU8 microgrooves using the Heidelberg MLA150 Maskless Aligner (Heidelberg Instruments, Heidelberg, Germany). Polydimethylsiloxane (PDMS) was prepared and cured on the micropatterned wafers according to manufacturer instructions (Sylgard Dow, Midland, Michigan). The PDMS molds were spin coated using the NiLo Scientific Spin coater 5 (NiLo Scientific, Ottowa, Canada) with a 5% PVA solution at 2000 rpm for 30 s. 0.8 mL of the 40% POCO solution (w/w in ethanol) was cured on the coated PDMS molds overnight at room temperature then 3-4 days at 80°C. The cured POCO and PDMS molds were submerged in DI water for 12-16 hrs at 60°C to dissolve the PVA sacrificial layer and remove the micropatterned POCO scaffold. The topography of the smooth and microgrooved scaffolds was evaluated using scanning electron microscopy (SEM). The scaffolds were coated with 7 nm osmium in an OPC-60A osmium coater (SPI supplies, West Chester, PA) and imaged with a Hitachi S-4800 microscope.

### In vitro cell biocompatibility

Biocompatibility of POC and POCO scaffolds was measured using an Alamar blue assay per the manufacturer’s protocol (Thermo Fisher). Leached scaffolds were incubated in complete cell medium (Lonza, Basel, Switzerland) for 2 days and treated with a 1 ug/mL fibronectin coating for 1 hr at room temperature. L929 fibroblast (Thermo Fisher) and human bone marrow derived mesenchymal stem cells (hMSCs) (Lonza) were seeded on the coated scaffolds at 20,000 cells/cm^2^. 24 hrs after seeding, the cell medium was replaced and with fresh medium and Alamar blue dye at a 1:10 dilution. The cells were incubated for 4 hrs at 37°C and the fluorescent signal of the Alamar blue and medium mixture was read at 590 nm on a Cytation 5 Cell Imaging Multi-Mode Reader (Agilent BioTek). The final fluorescence readings were adjusted from a blank and calculated as percent difference in reduction compared to a positive control and plotted. Prism9 was used to perform a one-way ANOVA.

Cell attachment and alignment were also evaluated with immunocytochemistry. The scaffolds were prepared as previously described and seeded with 10,000 MSCs/cm^2^. 7 days after seeding, the scaffolds were rinsed twice with PBS then fixed with a 4% PFA solution at room temperature for 30 min. The scaffolds were rinsed twice with PBS and permeabilized with 0.1% Triton-X100 at room temperature for 5 minutes. The scaffolds were rinsed twice with PBS again and blocked with 1% BSA at room temperature for 30 minutes. Next, scaffolds were stained with Phalloidin-iFlour (Abcam, Waltham, MA) and SYTOX green nucleic acid stain (Thermo Fisher) per the manufacturer’s protocol in the dark at room temperature for 20-30 min. Lastly, the scaffolds were washed three times with PBS for 5 min each. The final scaffolds were mounted on glass slides for imaging on a Cytation 5 Imaging Multi-Mode Reader (Agilent BioTek) or transferred to a glass bottom well plate for imaging on the Leica TCS SP5 confocal microscope (Leica Microscopes Inc, Deerfield, IL). Images were analyzed on ImageJ by identifying individual cells using segmentation and measuring aspect ratio and Feret angle. The results were plotted and analyzed using a Shapiro-Wilk test for normality and a two-tailed t-test in Prism9 software (n=30).

### Sample fixation for SEM imaging

Samples were fixed with 2.5% EM Grade gluteraldehyde and 2% paraformaldehyde in a 0.1M Sodium Cacodylate buffer solution and incubated overnight at 4C. The samples were washed with buffer and MilliQ water then incubated in 1% OsO_4_ for 1 hour at room temperature. Next, samples were rinsed with osmium and MilliQ water and dehydrated with a graded series of ethanol (30%, 50%, 70%, 85%, 95%, 100%). Samples then underwent critical point drying and were mounted on stubs for metal coating and imaging.

### Nude rat bladder augmentations

Microgrooved and smooth POCO scaffolds were evaluated for their potential to regenerate bladder tissue with and without seeded cells using an established nude rat bladder augmentation model.^22,23^ Prior to implantation, 1 cm^2^ scaffolds were incubated in complete MSC growth medium for 2 days and seeded with human bone marrow derived MSCs (Lonza) at a density of 15,000 cells/cm^2^. After 7-10 days, the medium was replaced with a 1:1 mixture of MSC growth medium and hematopoietic progenitor growth medium (Lonza) and human bone marrow derived CD34^+^ HSPCs (Lonza) were seeded on the scaffolds at 100,000 cells/scaffold. 24 hrs after seeding the CD34^+^ HSPCs, the scaffolds were evaluated for viability using NucBlue live cell stain (Thermo Fisher) and used for augmentation. Athymic nude rats (Charles River Laboratories, Skokie, IL) were used as a bladder augmentation model as previously described.^22,23^ Four experimental groups were evaluated: POCO and microgrooved (MG) POCO unseeded scaffolds, and POCO and microgrooved POCO scaffolds co-seeded with bone marrow-derived MSCs and CD34^+^ HSPCs. For all experimental groups, n=8 or 9 including a minimum of 4 female and 4 male rats.

### Urodynamics evaluation

Urodynamics were performed on anesthetized male and female rats prior to bladder augmentation surgery and on the day of euthanasia. Briefly, bladders of these animals were exposed through the abdomen and a 20-gauge needle was then inserted. The needle was then attached to a Pump 11 Elite Syringe Pump (Harvard Apparatus, Holliston, MA) and to a physiological pressure transducer (SP844, MEMSCAP, Isere, France). The transducer was attached to abridge amplifier (Model FE221; AD Instruments, Sydney, Australia), with constant pressure readings being displayed and calculated using LabChart 7.3 software (AD Instruments). Bladder capacity was also assessed at this time as sterile DPBS at a rate 150 to 200 μL per minute was infused until fluid leak was observed from the urethra. Voiding patterns were repeated three to five times per animal. Void frequency was calculated by determining the number of voiding events during total measurement time. Compliance was calculated by dividing change in bladder volume over the change in pressure. For all urodynamic data, results were plotted and analyzed using Prism9 software. Shapiro-Wilk tests were performed to assess for normal distribution. For void frequency and compliance, a paired t test or Wilcoxon test was performed to assess any difference in experimental groups pre- and 4 weeks post-augmentation. For bladder capacity, unpaired t tests were performed between the unseeded groups and the cell-seeded groups to assess any difference due to presence of microgrooves.

### Histological analysis and immunostaining

Bladder tissue was fixed in 10% buffered formalin phosphate (Thermo Fisher) before being dehydrated through a series of graded ethanol solutions (50%, 70%, 95%, 100%), and completed with xylene washes. After dehydration, samples were embedded in paraffin wax molds and sectioned onto glass slides using an RM2125 RT microtome (Leica Biosystems, Deer Park, IL) at a 5 µm thickness. Tissue sections were then stained with hematoxylin and eosin (H&E) or Masson’s trichrome. Furthermore, additional tissue sections underwent immunohistochemistry with the following primary antibodies; βIII tubulin, synaptophysin, pancytokeratin, uroplakin III, von Willebrand Factor, and CD31 with concentrations varying from 1:50–1:100 (Abcam, MA, USA). The slides were then mounted with Vectashield (Vector Laboratories, Newark, CA) containing 4′, 6-diamidino-2-phenylindole (DAPI) to extend fluorescence and visualize nuclei.

### Blood vessel quantification

The quantification of blood vessels was completed using a previously established protocol using 10-40x images of randomized regenerated tissue areas.^23^ Area of interest was determined by identifying the location of anastomosis between the native and regenerated tissue. Images were quantified in Adobe Photoshop CS3 (Adobe Systems Inc., San Jose, CA) following imaging with a Nikon Eclipse 50i microscope (Nikon Inc., Tokyo, Japan). Vasculature was manually quantified by applying the pen tool and recording pixel area from the image histogram tool. Quantification was completed by 10 blinded participants. Vascular quantification data are represented as the mean number of vessels per square millimeter, mean percent vasculature (means ± SE), and average vessel size. Data was plotted using Prism9 software and a Shapiro-Wilk test was performed to assess normal distribution of the data. Next, an unpaired t test or Kolmogorov-Smirnov test was performed between unseeded and cell-seeded groups to determine statistically significant differences between groups. The total number of vessels and corresponding size for each experimental group were plotted in histograms using Python to visualize the distribution of vessel size (Python code provided on request).

### Muscle quantification

Muscle quantification was achieved using images of the regenerated tissue area. The regenerated tissue area was determined by identifying the location of anastomosis. Ten images spanning the regenerated tissue were assessed in Adobe Photoshop first by removing vasculature and urothelium with the eraser tool. Color range was then digitally selected for red or blue pixels for the whole image and artifacts were removed. Those selected pixels were quantified using the image histogram tool and a muscle to collagen ratio was established. Results were assessed for normal distribution with a Shapiro-Wilk test, and an unpaired t test or Kolmogorov-Smirnov test was performed to determine statistically significant differences between unseeded and cell-seeded. Data was plotted and analyzed using Prism9 software.

### Urothelium quantification

Urothelium quantification was performed on Masson’s trichrome-stained tissue sections. Two, 10x images were obtained spanning the length of the anastomosis ends and the regenerated bladder tissue. Using Spot Advanced Imaging software (Diagnostic Instruments, Sterling Heights, MI), ten measurements were obtained by measuring the distance from the lamina propria to the urothelium lumen. Shapiro-Wilk tests and unpaired t tests or Kolmogorov-Smirnov tests were performed to assess normality and statistically significant differences between unseeded and cell-seeded groups. Data was plotted and analyzed using Prism9 software.

### Peripheral nerve quantification

Following the staining of bladder tissue with antineuronal antibody, βIII tubulin, peripheral nerve quantification was performed using Image Spot Software (Diagnostic Instruments). With measurement settings calibrated to 40x binning none, the measurement tool was used to record the length of nerve elements. Neural innervation was quantified by measuring the distance between the anastomosis of the native-regenerated tissue border and the farthest instance of βIII tubulin^+^ staining for each animal. Quantification was completed by 3 blinded participants and is represented as the mean length of nerves in micrometers. Data was plotted using Prism9 software and Shapiro-Wilk tests and unpaired t tests or Kolmogorov-Smirnov tests were performed to assess normality and statistically significant differences between unseeded and cell-seeded groups.

## Results

### POCO scaffold mechanical properties are similar to native bladder tissue

Both the ^1^H-NMR and FTIR spectra confirmed the expected chemical structure of the polymer **(Fig 2C-D)**. In particular, the presence of octanol in POCO was verified from the peak at 0.86 ppm in the ^1^H-NMR spectra. The integral of this peak was normalized and compared to the integral of the citric acid group (2.77 ppm) to calculate the percent octanol content. This was found to be 18%, which is comparable to the molar ratio of octanol used in POCO synthesis.

**Figure 1:**
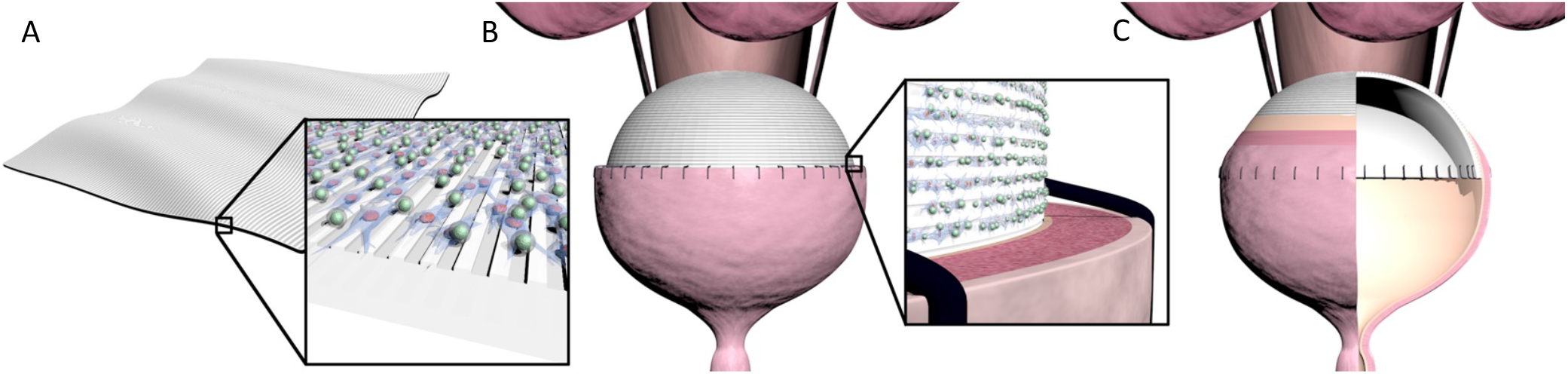
Schematic of cell-seeded, microgrooved POCO scaffold promoting bladder regeneration following augmentation. (A) Microgrooved POCO scaffold seeded with bone marrow-derived mesenchymal stem cells (MSCs) and CD34^+^ hematopoietic stem/progenitor cells (HSPCs). (B) Bladder augmented with cell-seeded, microgrooved POCO scaffold (C) Regenerating bladder with microgrooved POCO scaffold.

**Figure 2:**
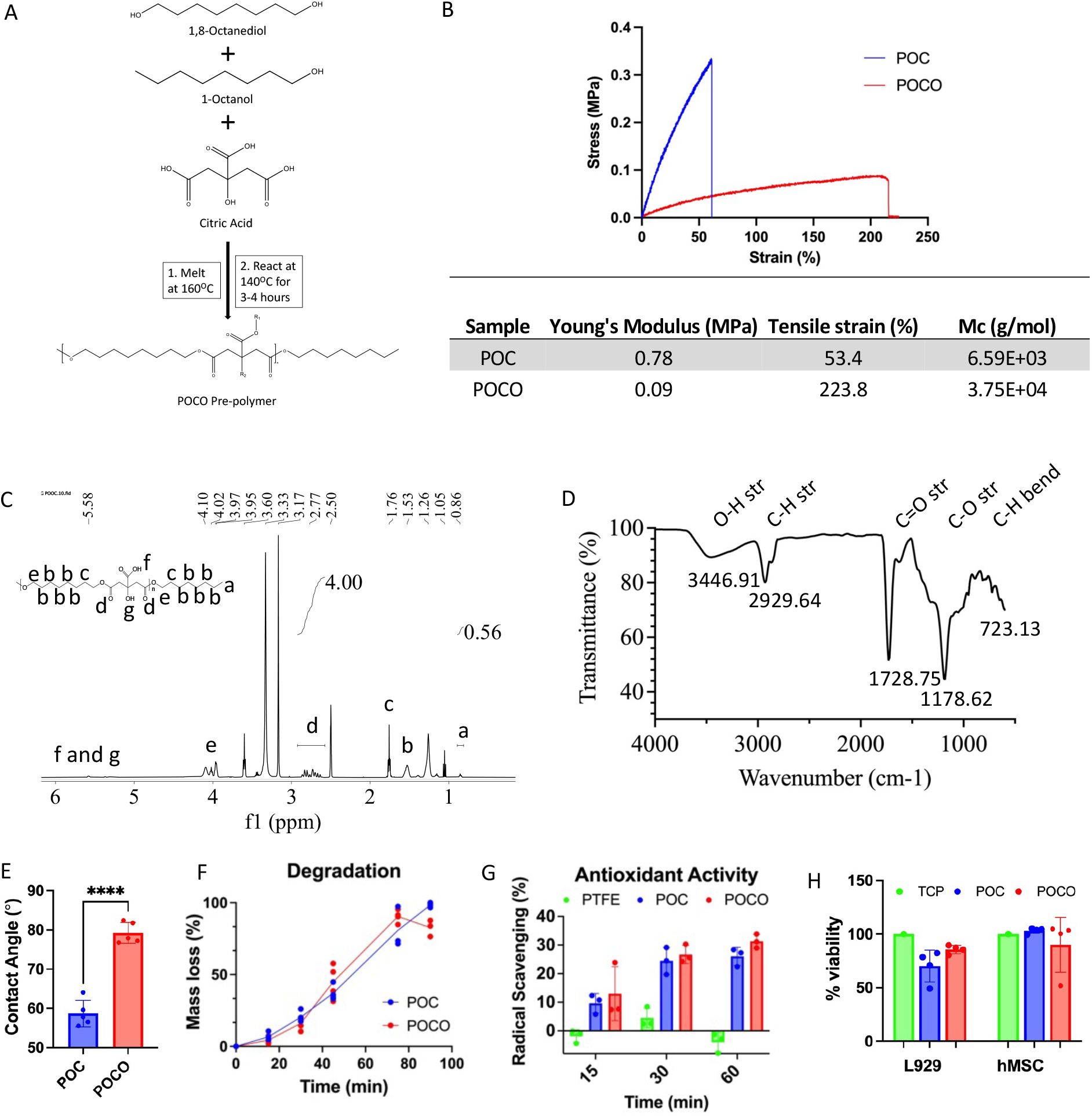
POCO and POC characterization. (A) Schematic of POCO synthesis. (B) Representative stress-strain curve and table with mechanical properties of POC and POCO (C) ^1^H-NMR spectra of POCO with calculated integrals. (D) FTIR spectra of POCO. (E) Contact angle of POC and POCO (^****^ p< 0.001). (F) Accelerated degradation curve of POC and POCO in 0.1 M NaOH. (G) Antioxidant activity of POC and POCO compared to PTFE (negative control) (H) Biocompatibility of POC and POCO with L929 fibroblasts and hMSCs.

As expected, molecular weight between crosslinks was significantly higher for POCO compared to POC **(Fig 2B)**, and crosslinking density of POC scaffolds was found to be significantly higher than that of POCO, even when POCO was cured for longer times **(Supp Fig 2)**. The Young’s Modulus of the POCO scaffolds is one-fifth that of POC, and within the normal range for healthy rat bladder tissue at approximately 0.2 MPa **(Fig 2B)**.^26^ The tensile strain of POCO scaffolds is about 4 times that of POC, reaching over 200% elongation before break **(Fig 2B)**. These properties are critical for bladder tissue regeneration because the scaffold must be able to mimic the native bladder tissue and withstand cyclic expansion and contraction. The addition of octanol also affected contact angle of the scaffolds. The methyl group from the octanol increases hydrophobicity of POCO compared to POC **(Fig 2E)**.

The presence of citrate bonds in POCO and POC confer the materials with antioxidant properties, which can be beneficial in bladder applications to prevent stone formation.^27^ POCO and POC were found to have comparable antioxidant activity **(Fig 2G)**. The accelerated degradation curves of POCO and POC are also similar with both scaffolds reaching about 50% degradation at 45 min and 100% degradation at 90 min at room temperature in 0.1 M NaOH solution **(Fig 2F)**. Lastly, the biocompatibility of POCO and POC was compared using mouse L929 fibroblast and hMSCs. For both cells types and scaffolds, the cell viability remained around 100% with no significant difference compared to a tissue culture plasticware control after 24 hrs **(Fig 2H)**.

### In vitro characterization of microgrooved POCO scaffolds

After validating the chemical and mechanical properties of POCO, microgrooved POCO scaffolds were fabricated to promote cell alignment. The structure of the microgrooves was confirmed with SEM imaging **(Fig 3B)**. All microgrooved scaffolds appeared to have uniform grooves 50 μm apart and 10 μm deep. hMSC attachment and alignment on the microgrooved POCO scaffolds were visually and quantitatively assessed using fluorescence microscopy **(Fig 3C-E)**. Visual inspection of the fluorescence images suggested that the hMSCs on the microgrooved POCO had a more elongated phenotype and followed the direction of the microgrooves. Cell morphology in the microgrooves was quantified by measuring aspect ratio of the hMSCs, which was significantly higher on the microgrooved POCO compared to TCP **(Fig 3D)** (p< 0.01). Similarly, cell orientation was quantified through measuring Feret angle. While the Feret angle for the hMSCs on the microgrooved surface varied from 120°-165°, Feret angle on the TCP varied from 0°-180°. The smaller variance in Feret angle on the Microgrooved POCO suggests that cells orient along the grooves. These results confirm greater cell elongation and alignment in the microgrooves.

**Figure 3:**
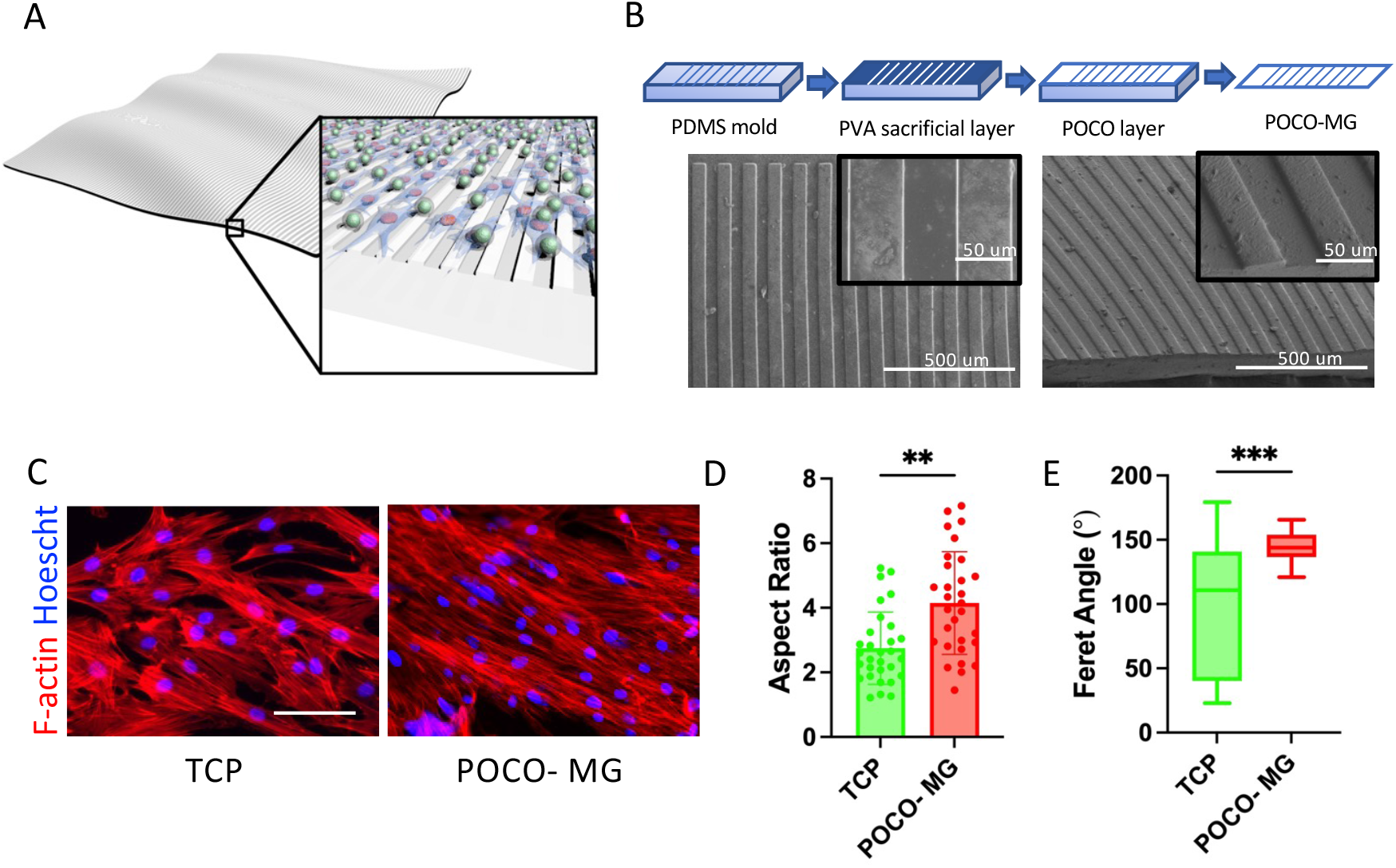
*In vitro* characterization of POCO. (A) Microgrooved (MG) POCO seeded with MSCs and HSPCs (B) Schematic of MG scaffold fabrication and SEM images of MG POCO (C) Fluorescence images of hMSCs on a tissue culture plate (TCP) and MG POCO (D) Aspect ratio of hMSCs on TCP vs MG POCO (n=30) (^**^ p< 0.01) and (E) Feret angle of hMSCs on TCP vs. MG POCO (n=30) (^***^ p< 0.001).

### POCO scaffolds restore normal urodynamics in augmented bladders

Urodynamic studies (UDS) were used to compare voiding patterns and bladder function in a previously established nude rat bladder augmentation model pre- and 4 weeks post-augmentation. From the UDS tracings pre- and post-augmentation, no change in frequency or magnitude of voids was observed in either male or female rats **(Fig 4A)**. Volumetric capacity pre- and 4 weeks post-surgery was measured to determine if the scaffolds successfully augmented the bladder by increasing capacity **(Fig 4B)**. Only the Smooth-cell seeded and Microgrooved-cell seeded groups demonstrated increased capacity at 4-weeks post augmentation. On average, these experimental groups demonstrated 30% increased volume compared to the original capacity, while the Smooth and Microgrooved groups demonstrated approximately 10% decrease from the original capacity. Void frequency pre- and post-augmentation also differed between the unseeded and cell-seeded experimental groups. On average, void frequency of the unseeded groups increased by over 100% (1 void/min to 2.2-2.3 voids/min post-augmentation, p<0.001) while void frequency of the cell-seeded groups increased by 15-20% (1-1.2 voids/min to 1.3-1.4 voids/min, p<0.05) post-augmentation **(Fig 4C)**. When comparing compliance, the Microgrooved and Smooth-cell seeded groups demonstrated significant changes post-augmentation **(Fig 4D)**. The majority of the Microgrooved group experienced a decrease in compliance (p< 0.05), while the Smooth-cell seeded group experienced an increase in compliance (p< 0.01). Decreased compliance can be indicative of bladder tissue that is not completely malleable, but despite any changes post-augmentation all the experimental groups demonstrated compliance measurements that were within the normal range.

**Figure 4:**
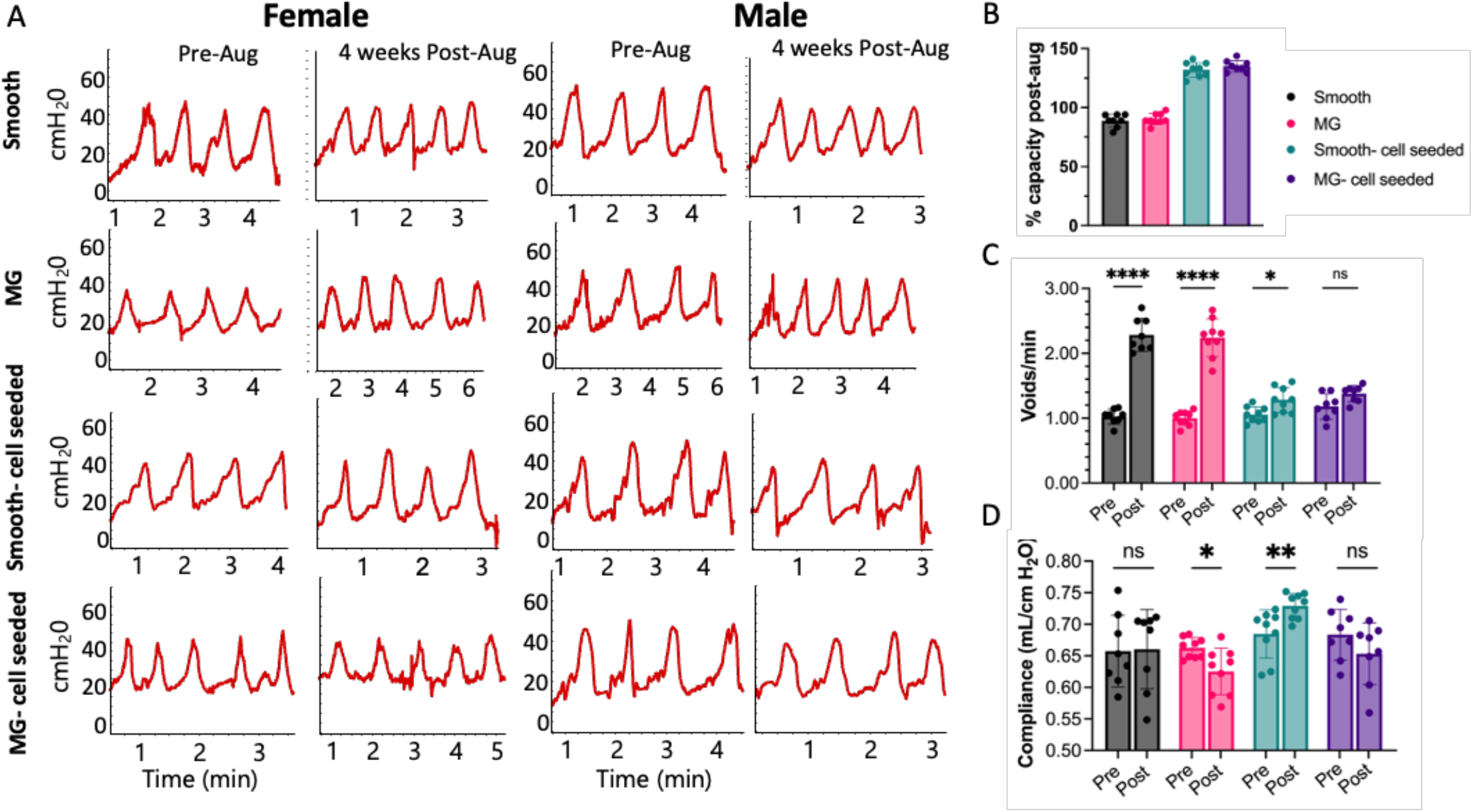
Urodynamics pre- and 4-weeks post-augmentation. (A) Representative urodynamic tracings for male and female rats (B) capacity pre- and post-augmentation (n=8) (C) void frequency pre- and post-augmentation (n=8) (^****^ p< 0.001, ^*^ p< 0.05) (D) compliance measurements pre- and post-augmentation (n=8) (^*^ p< 0.05, ^**^ p< 0.01).

### Microgrooved scaffolds enhance urothelium and peripheral nerve regeneration

Regenerated bladder tissue was assessed for muscle, urothelium, and peripheral nerve regeneration. Trichrome staining was used to measure muscle: collagen ratio, determine presence of fibrotic tissue, and observe collagen alignment **(Fig 5A)**. Here, we observed that regenerated tissue from the cell-seeded groups had muscle: collagen ratios similar to that of native tissue at approximately 0.6, which is twice that of the unseeded groups. Between the Smooth-cell seeded and MG-cell seeded groups, muscle: collagen ratio increased from an average of 0.53 to 0.61 (p<0.01). Urothelium regeneration was evaluated by measuring urothelium thickness and expression of markers pancytokeratin and uroplakin III (UPIII) **(Fig 5B)**. The presence of microgrooves induced regeneration of a thicker and more uniform urothelium layer. The addition of cells also increased average urothelium thickness, with the cell-seeded groups reaching an average urothelium thickness that is comparable to native tissue at approximately 51μm **(Fig 5B)**. When comparing the urothelium in the unseeded groups, the addition of microgrooves allowed for uniform expression of UPIII across the regenerated urothelium **(Fig 5B)**. Lastly, nerve regeneration was assessed via staining for peripheral nerve markers synaptophysin and β-III tubulin **(Fig 5C)**. The presence of cells dramatically increased both average nerve length and nerve infiltration. Between the Smooth-cell seeded and MG-cell seeded groups, average peripheral nerve average length decreased from 40.5 μm to 37.3 μm (p<0.05), but all experimental groups had a lower average nerve length comapred to native tissue. The microgrooves also appeared to have an effect on the ingrowth of nerves into the regenerated tissue: only 62.5% of the Smooth group had apparent nerve regeneration compared to 88.9% of the microgrooved group.

**Figure 5:**
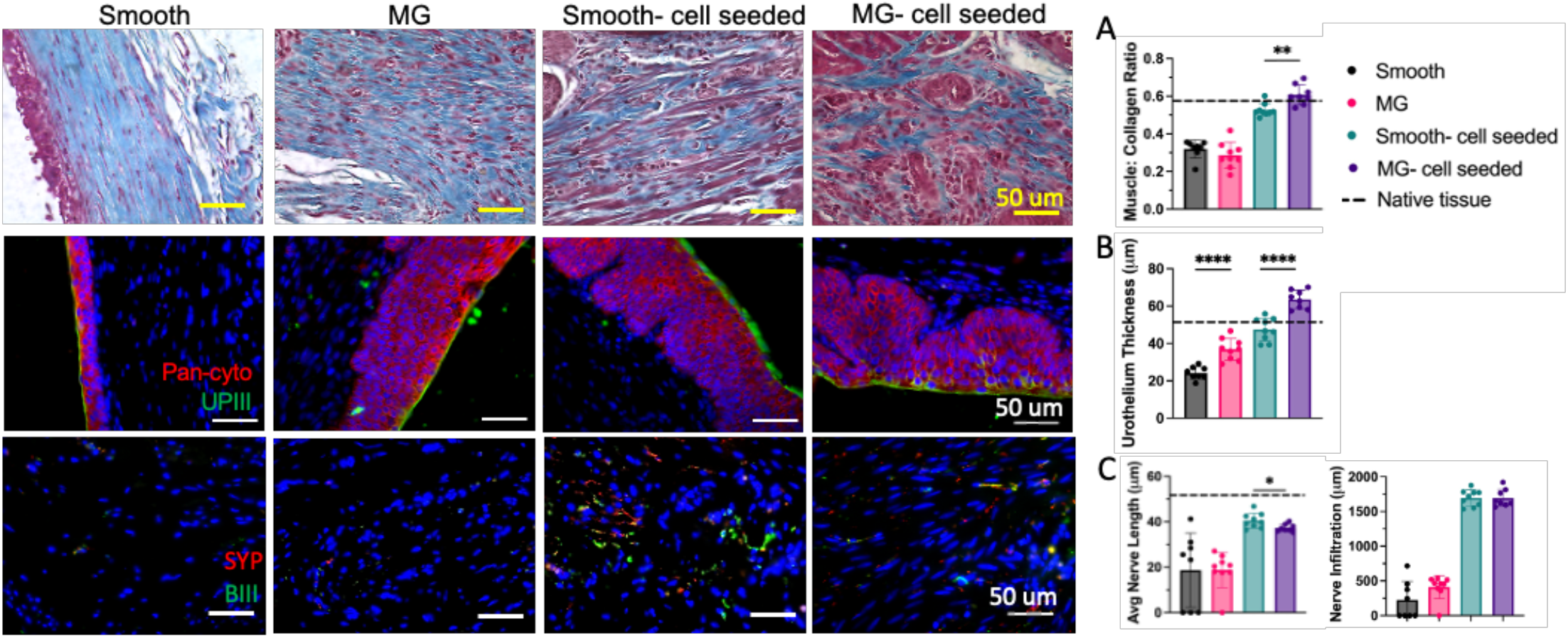
Muscle, urothelium, and nerve regeneration. (A) Representative trichrome staining of regenerated bladder tissue and corresponding quantification of muscle to collagen ratio (n=8) (^**^ p<0.01). (B) Imrnunostaining for urothelium markers Pan cytokeratin (Pan-cyto) (red) and uroplakin III (UPII) (green) with corresponding quantification of urothelium thickness (n=8) (^****^ p<0.0001). (C) Immunostaining for nerve markers synaptophysin (SYP) (red) and beta-III tubulin (BIii) with quantification of nerve migration into the regenerated tissue and average nerve length (n=8) (^*^ p< 0.05).

### Microgrooved MSC/CD34^+^ scaffolds support increased neovascularization

To evaluate revascularization, we stained regenerated tissue for vasculature markers vonWillebrand Factor (vWF) and CD31 **(Fig 6A)**. Trichrome staining was used to measure surface area of vasculature in the regenerated tissue as well as the number of blood vessels. The presence of vessels was confirmed with positive staining of vWF and CD31. Here the cell-seeded groups showed 2-3 times the number of vessels in the regenerated tissue when compared to the unseeded groups **(Fig 6B)**. Interestingly, the MG-cell seeded group had a lower relative surface area of vasculature in the regenerated tissue compared to the Smooth-cell seeded group (p<0.05) and a higher number of vessels (p<0.0001) **(Fig 6B-C)**. All of the observed blood vessels in each experimental group were sorted based on diameter and plotted in histograms to compare vessel size distributions for cell-free and cell-seeded scaffolds **(Fig 6D)**. The vast majority of the blood vessels in the Microgrooved cell-seeded group ranged from 50-100 μm as compared to the Smooth cell-seeded group, which had a more even vessel size distribution. This trend was confirmed by measuring average vessel size across all experimental groups, which showed that the presence of microgrooves in the cell seeded groups decreased average vessel size by about 100 μm (p<0.0001) **(Fig 6E)**.

**Figure 6:**
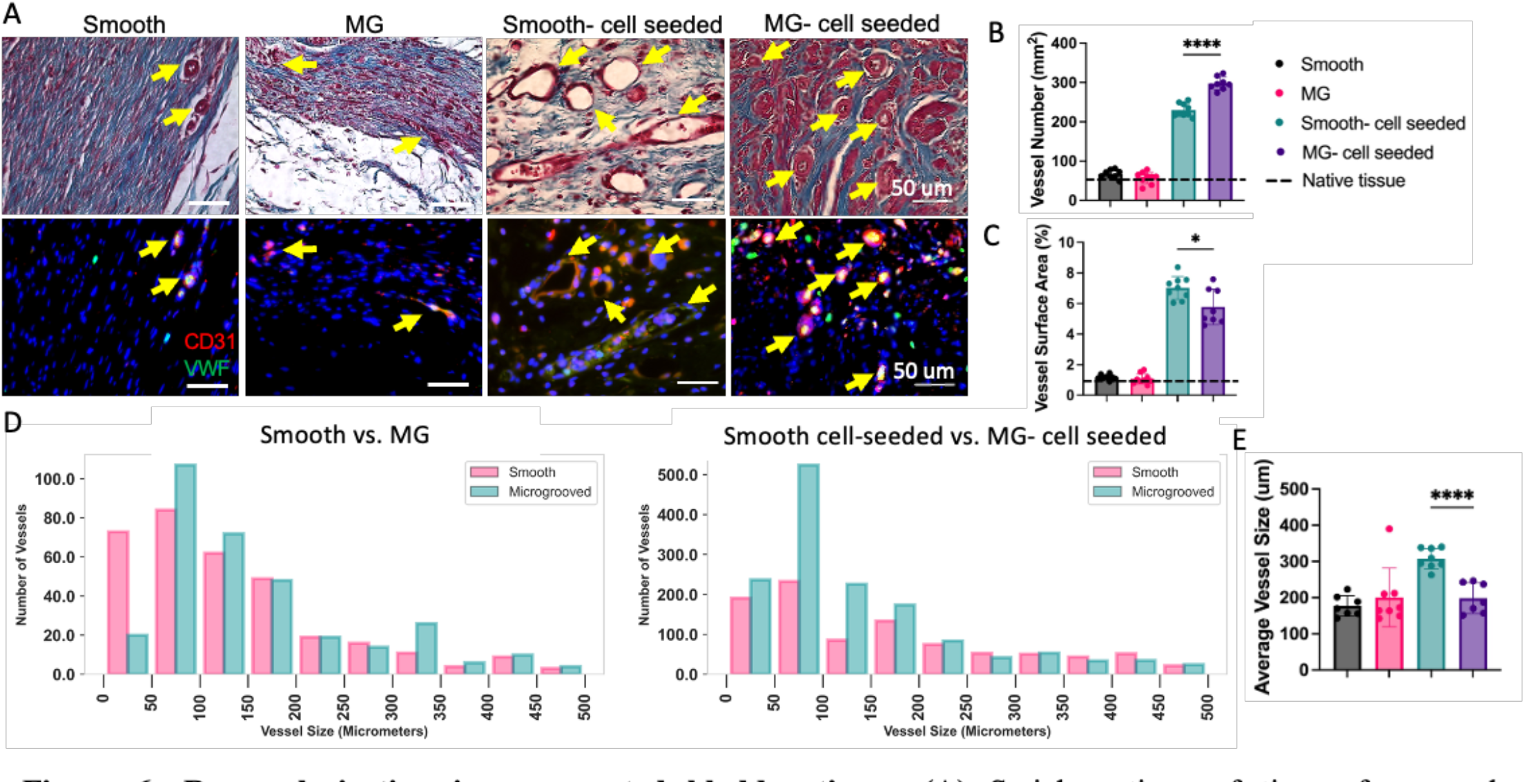
Revascularization in regenerated bladder tissue. (A) Serial sections of tissue for vascularquantification (rows 1-2). Yellow arrows indicate blood vessels in trichrome stains and immunostaining for vonWillebrand Factor (vWF) (green) and CD31 (red). (B) Number of vessels (n=8) (^****^ p<0.0001) (C) Surface area of vasculature (n=8) (^*^ p< 0.05). (D) Histograms of vessel size distribution represented as cumulative vessels per experimental group in animals that received unseeded (left) and seeded (right) scaffolds, and (E) overall average vessel sizes (n=8) (^****^p< 0.0001).

## Discussion

The ideal graft for bladder tissue engineering should provide physical and biological cues to promote efficient regeneration of all bladder wall components while supporting bladder function throughout recovery from augmentation. Previous work on the development of citrate-based elastomer scaffolds for bladder tissue engineering showed promising results with robust regeneration of the bladder wall in rat and baboon augmentation models.^21,22,27–29^ However, these scaffolds required further optimization to better promote vascular and peripheral nerve regeneration, which are critical for healthy tissue formation and function. For this study, we evaluated the effect of microtopography engineering on bladder tissue regeneration; specifically, to determine if the presence of microgrooves would induce cell alignment and migration for improved tissue function and tissue quality.

*In vitro* validation of the microgrooved POCO scaffolds confirmed that the material mimics mechanical properties of native bladder tissue and causes a distinct phenotypic change to the seeded cells. hMSCs seeded on the microgrooved POCO scaffolds demonstrated greater elongation and uniform alignment in the direction of the microgrooves. These scaffolds were then tested in an established rat bladder augmentation model to determine if the morphology change to the seeded cells would alter the subsequent tissue regeneration profile.

Bladder function was first evaluated with urodynamic testing pre- and post-augmentation with the cell seeded scaffolds. Urodynamic testing is common practice in a clinical setting as it tests the ability of the bladder to store and expel urine through measures such as capacity, void frequency, and compliance. Capacity is defined as the maximum bladder volume and quantifies the bladder’s storage function. In the cell seeded groups, capacity increased by about 30% at 4 weeks post-augmentation, while capacity of the unseeded groups remained relatively similar pre- and post-augmentation. It has been well-established that the presence of stem/progenitor cells supports the regeneration of a robust bladder wall rather than fibrous tissue formation.^30,31^ Therefore, the increased capacity in the cell seeded groups suggests successful augmentation of the bladder and implies growth of new bladder tissue. These results are consistent with the changes to void frequency pre- and post-augmentation. For unseeded groups, void frequency doubled post-augmentation while the cell seeded groups only had a modest increase in void frequency. This difference may be attributed to the increase in capacity and presence of healthier bladder tissue for the cell seeded groups. Similar results were observed for changes in compliance post-augmentation. Compliance measures the ability of the bladder to maintain low pressure during filling. Non-compliant tissue significantly hinders function as it limits the expansion and contraction cycles, and fibrotic tissue in particular has been shown to be less compliant than healthy bladder tissue.^32^ Only the Microgrooved group showed a slight decrease in compliance post-augmentation, but all animals maintained normal compliance over the course of the study. Overall, it appears that microtopography did not have a significant effect in terms of bladder function; however, our results suggest that a cell-free POCO scaffold supports normal function and limits scar tissue formation, representing an important step towards development of an alternative graft material for BAE.

Histological evaluation of the regenerated bladder wall was also conducted to quantitatively assess any differences in the new tissue that arose from the presence of the microgrooves. The first component of the bladder wall to regenerate is the urothelium, which plays a critical role in creating a water-tight barrier for storage of urine from the rest of the body.^33–35^ Surprisingly, we found that the presence of microgrooves enhanced urothelium regeneration independent of the presence of cells. The uniformity and thickness of the urothelium significantly increased on microgrooved scaffolds, suggesting that the presence of microgrooves alone is sufficient to promote migration of native urothelial cells into the graft and form a complete water-tight barrier at 4-weeks post-augmentation. Furthermore, the microgrooved scaffolds promoted rapid development as indicated by urothelium thickness and expression of UPIII, a mature urothelium marker representing formation of an impermeable barrier in the bladder.^33^ The urothelium allows the subsequent components of the bladder wall to regenerate and plays a critical role in sensing physical and chemical stresses on the bladder and transducing these signals to the rest of the bladder cell types. Previous work has shown that the urothelium also influences smooth muscle development and patterning during regeneration.^36^ The ability of the urothelium to not only protect but modulate bladder activity suggests that promoting rapid and robust regeneration of the urothelium via microtopography engineering can influence subsequent success of the overall bladder recovery post-augmentation. Furthermore, these results are consistent with those of recent work in regenerating epithelial layers in other tissues (e.g. vascular networks and skin) with micropatterning.^37–39^ In these studies, grooved patterns in particular were observed to promote increased epithelial cell migration and alignment. As such, incorporation of microtopography engineering into POCO scaffolds shows promise to accelerate and improve bladder regeneration.

Regeneration of the detrusor muscle is also necessary to recover bladder function post-augmentation; specifically, the coordinated expansion and contraction to store and expel urine. Successful regrowth of the detrusor muscle is an ongoing challenge as scaffolds are prone to formation of fibrous tissue, which lacks the mechanical properties and contractility of native smooth muscle. Herein we observed that bladders augmented with cell-seeded scaffolds displayed significantly higher muscle to collagen ratios, closer to the ratio observed in native tissue. This effect may be attributed to paracrine effects and/or differentiation of the seeded cells. MSCs in particular have similar phenotypes to bladder smooth muscle cells and were previously observed to integrate into smooth muscle bundles.^12^ On average, the Microgrooved-cell seeded group demonstrated slightly higher muscle:collagen ratios compared to the Smooth-cell seeded group. This finding may be due to the alignment of smooth muscle cells or fibers in the presence of microgrooves. However, further investigation is necessary to confirm this hypothesis.

The remaining components of the bladder wall that are essential for complete regeneration are peripheral nerve and vascular networks. Regeneration of these structures has historically been difficult to achieve in a bladder augmentation setting, but they are critical to sustain growth of the bladder wall and restore function.^9,10^ Previous work with cell-seeded citrate-based elastomer scaffolds demonstrated evidence of robust revascularization and peripheral nerve ingrowth.^29^ The grafted cells were observed to integrate into neovasculature and provide increased blood flow to support peripheral nerve growth. As anticipated, augmentations with cell-seeded scaffolds in this study showed a significant increase in revascularization and nerve regeneration compared to that of the cell-free scaffolds, and the extent of vascular and nerve regeneration is analogous to previous publications.^22,29^ Furthermore, the presence of microgrooves appeared to induce ingrowth of nerves in the unseeded Microgrooved group. Approximately 60% of the smooth group had observed nerve regeneration compared to about 90% of the Microgrooved group. These results are promising for future studies developing scaffolds that can promote rapid peripheral nerve regeneration in the bladder. The microtopography also altered the number of blood vessels in the regenerated tissue. The vessel number increased by about 30% between the cell seeded groups, and the majority of the vessels observed in the Microgrooved-cell seeded group were 50-100 μm in diameter. Therefore, the microgrooves appear to encourage alignment to better promote revascularization. To the best of our knowledge, this is the first report to examine the role of scaffold microtopography and cell contact guidance in bladder tissue regeneration. Optimizing the microtopography patterns will be necessary to assess the impact of this approach. Therefore, this study provides novel insight on the application of microtopography engineering to direct regeneration of the bladder post-augmentation.

Despite the promising results using microtopography engineering to improve scaffold performance for bladder regeneration, there are limitations. First, the study was conducted in an immunocompromised, small animal model. To better predict how this system would perform in human patients, the study should be conducted in a large, immune competent animal model with similar micturition behavior. Furthermore, the duration of the study was only 4 weeks, which may not be long enough to observe optimal peripheral nerve regeneration and revascularization in a rat model. Previous studies have shown that recovery of robust nerve networks is not observed even after 10 weeks post-augmentation in rat models; therefore, studies that occur over longer time periods may provide more insight into the effect of microtopography on nerve regeneration.^27,29^ Another potential shortcoming of this study is the chosen size and pattern of the microgrooves. The width, depth, and pitch were observed to promote alignment of the seeded MSCs; however, a variety of microtopographies should also be explored to examine the effect of the microgroove dimensions and pattern on cell alignment and migration in the bladder environment. A concentric circle pattern may be of interest to study given that it could mimic the alignment of muscles in the bladder wall as well as grooves that could mimic native vasculature patterns. Nonetheless, this work presents a new avenue of research in the bladder regenerative engineering space that can be explored to develop a better alternative graft for patients requiring bladder augmentations. Given that the role of contact guidance in bladder tissue engineering is largely unknown, this study illustrates how microtopography engineering can be implemented to influence bladder tissue regeneration and potentially, the regeneration of other organs.

## Data availability

Data available on request from the authors

## Acknowledgements

This research was supported by Center for Advanced Regenerative Engineering and the Department of Urology at the Feinberg School of Medicine at Northwestern University, the Division of Urology at Lurie Children’s Hospital of Chicago, NIH grant R01EB026575 to A.K.S. and G.A.A, NIH grant R01DK109539 to A.K.S., and T32 grant (EB031527) to G.A.A

*This work made use of the NUFAB facility of Northwestern University’s NUANCE Center, which has received support from the SHyNE Resource (NSF ECCS-2025633), the IIN, and Northwestern’s MRSEC program (NSF DMR-1720139). This work made use of the IMSERC NMR facility at Northwestern University, which has received support from the Soft and Hybrid Nanotechnology Experimental (SHyNE) Resource (NSF ECCS-2025633), Int. Institute of Nanotechnology, and Northwestern University*.

*This work made use of the Keck-II and BioCryo facilities of Northwestern University’s NUANCE Center, which has received support from the SHyNE Resource (NSF ECCS-2025633), the IIN, and Northwestern’s MRSEC program (NSF DMR-1720139)*.

*Microscopy was performed at the Biological Imaging Facility at Northwestern University (RRID:SCR_017767), graciously supported by the Chemistry for Life Processes Institute, the NU Office for Research, the Department of Molecular Biosciences and the Rice Foundation*.

*The authors would like to thank Mark Seniw at the Simpson Querrey Institute at Northwestern University for creating the content found in Figure 1*.

## Supplementary Figures

**Supplementary Figure 1:**
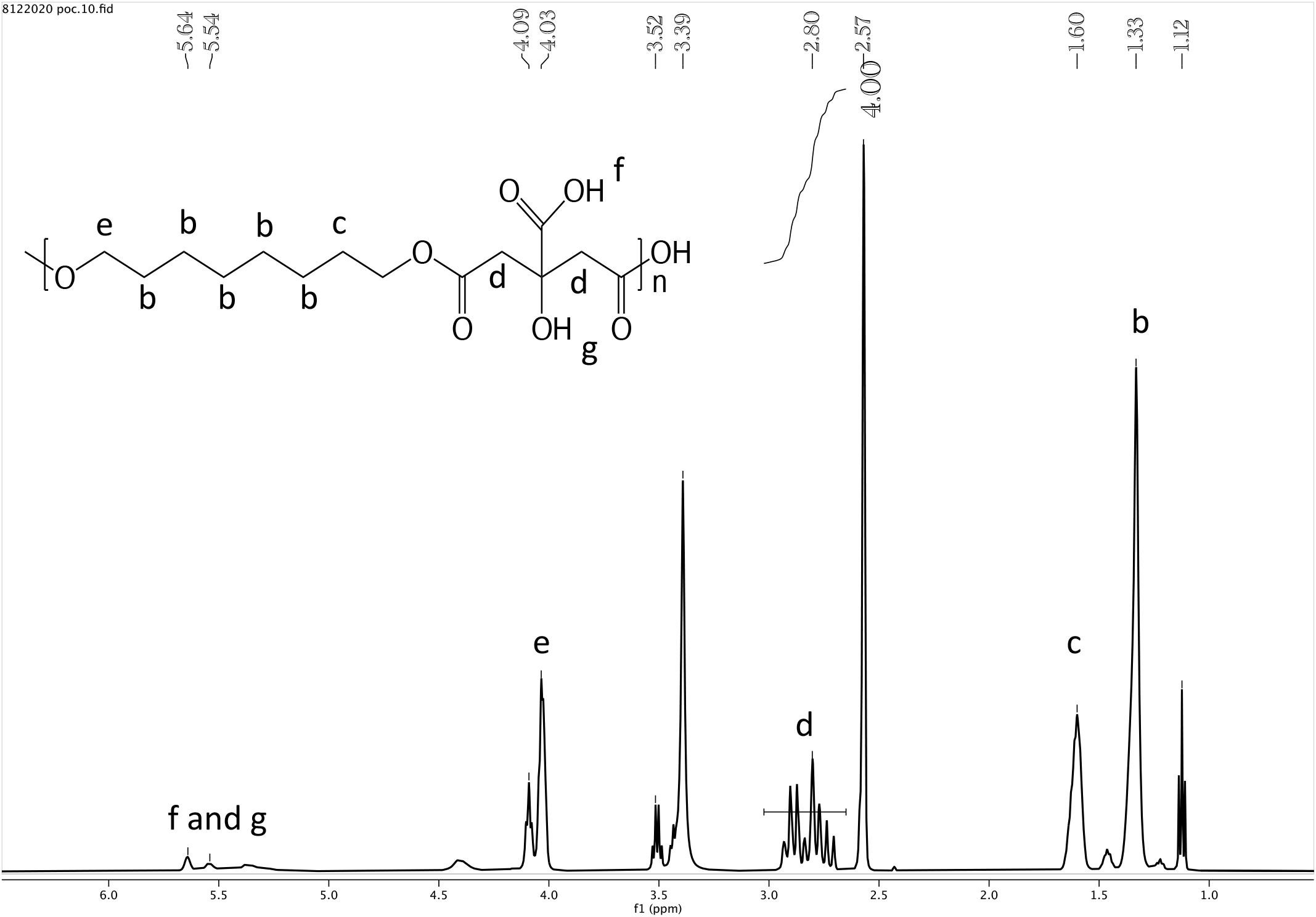
^1^H-NMR spectrum of POC. Compared to POCO, the ^1^H-NMR spectrum of POC has no peak at 0.86 ppm, which indicates presence of the methyl group from octanol.

**Supplementary Figure 2:**
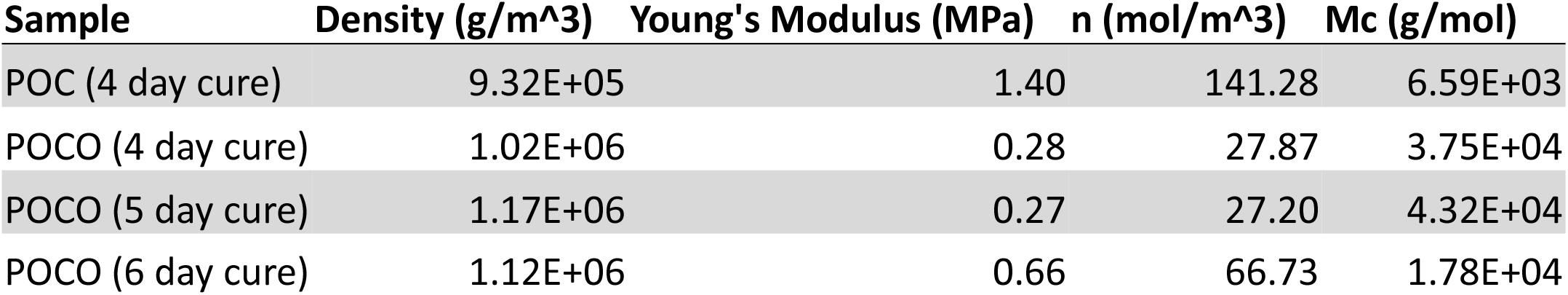
Comparison of density, Young’s Modulus, molecular weight between crosslinks (n), and crosslinking density (M_c_) between POC and POCO under different curing conditions.

**Supplementary Figure 3:**
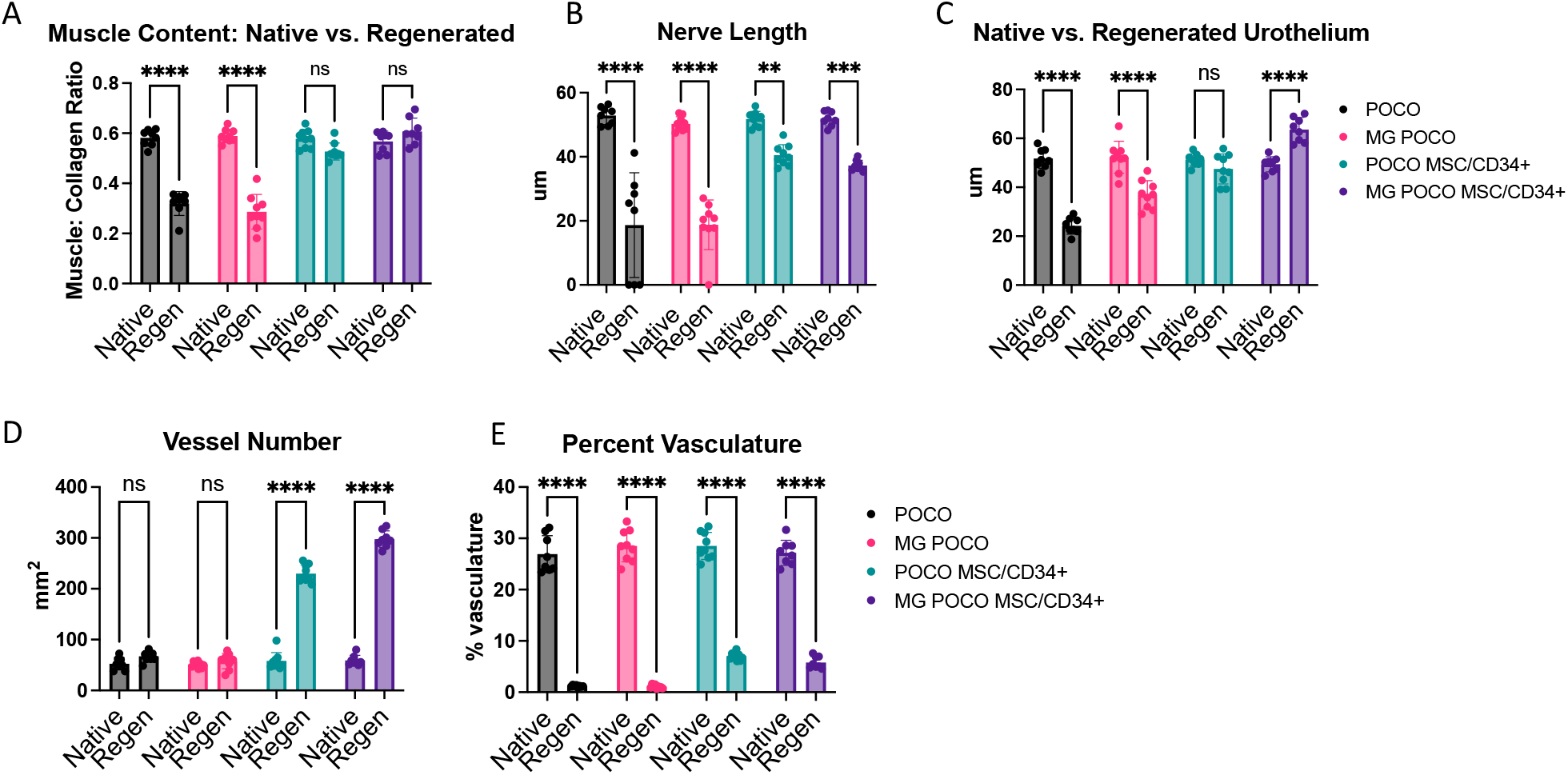
Comparison of native and regenerated tissue. (A) Muscle: collagen ratio (^****^p < 0.001) (B) Average nerve length (^**^ p< 0.01, ^***^ p< 0.005, ^****^p < 0.001) (C) Average urothelium thickness (^****^p < 0.001) (D) Average number of blood vessels (^****^p < 0.001) (E) Surface area of vasculature (^****^p < 0.001)

**Supplementary Figure 4:**
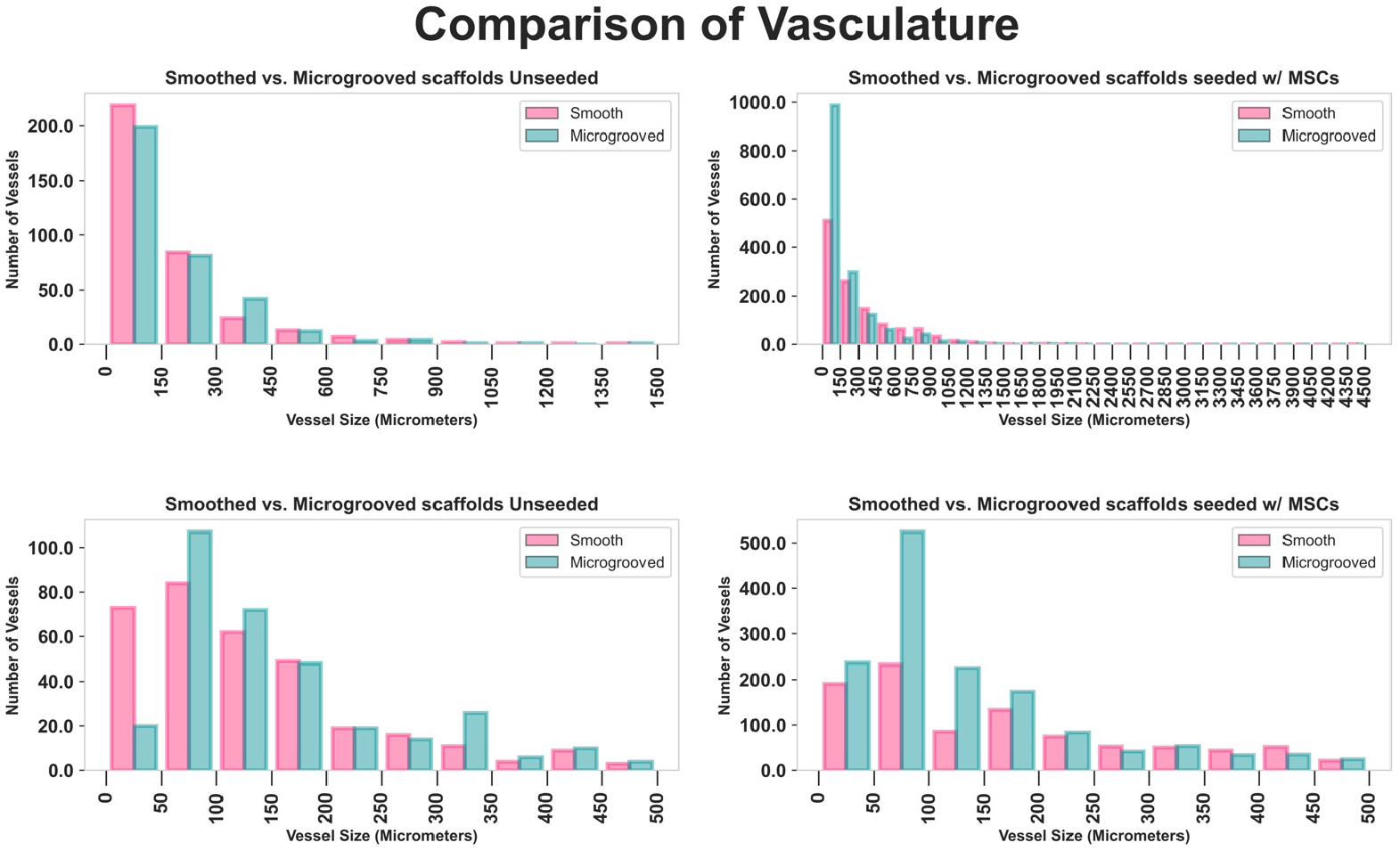
Complete histogram of vessel size distribution in regenerated tissue represented as total vessels per experimental group. POCO and MG POCO (left) with full vessel size spectrum (top) and 0-500 um range (bottom). POCO MSC/CD34^+^ and MG MSC/CD34^+^ (right) with full vessel size spectrum (top) and 0-500 um range (bottom).

**Supplementary Figure 5:**
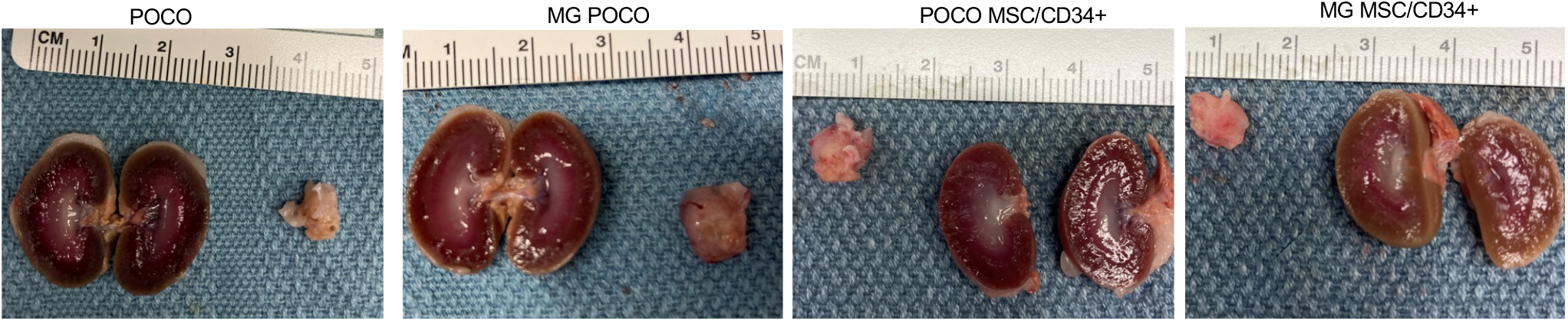
Gross histology of rat kidneys and bladder at 4 weeks post-augmentation

**Supplementary Figure 6:**
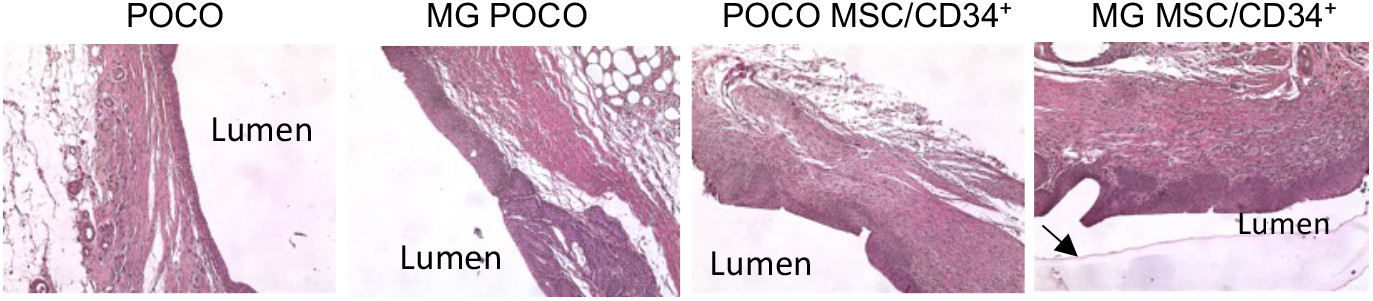
Representative H&E staining of regenerated tissue at 4 weeks post-augmentation. Black arrow indicates remaining POCO scaffold facing the lumen of the bladder wall.

